# The 10 bits/s bottleneck as error-correcting redundancy: an information-theoretic theory of cognitive reserve

**DOI:** 10.64898/2026.03.24.713899

**Authors:** Don Yin

## Abstract

Among individuals with equivalent Alzheimer’s pathology, cognitive outcomes can diverge by decades, a phenomenon termed cognitive reserve that remains descriptive after thirty years of research. We propose that the ∼10^9^-to-10 bits/s gap between sensory input and behavioral output functions as error-correcting redundancy in the sense of Shannon’s channel coding theorem. Progressive neuronal loss maps to symbol erasure in a redundant code, and the critical damage fraction at which cognition fails is *d*_*c*_ = 1 − *k/n*, where *k* ≈ 10 bits/s is the behavioral channel requirement and *n* is the effective number of coding units. We evaluate this threshold across three channel models (binary erasure, Gaussian, and Erdős–Rényi percolation) and show that all produce a sharp phase transition from reliable to unreliable decoding. The framework makes four testable predictions: (i) *d*_*c*_ scales with the measurable redundancy ratio *ρ* = *n/k*, which accounts for clinical heterogeneity; (ii) information-theoretic redundancy from resting-state fMRI should predict time-to-conversion beyond structural atrophy; (iii) the decline trajectory near *d*_*c*_ is sharp, consistent with the “cognitive cliff”; and (iv) motor circuits, operating at higher bandwidth, have lower reserve than cognitive circuits.

**Significance Statement:** Cognitive reserve (why some brains resist dementia pathology better than others) has been described for thirty years but never given a quantitative, information-theoretic foundation. We propose that the roughly hundred-million-fold gap between sensory input (∼10^9^ bits/s) and behavioral output (∼10 bits/s) functions as error-correcting redundancy in the Shannon coding-theoretic sense. This yields a closed-form critical damage threshold, *d*_*c*_ = 1 − *k/n*, below which cognitive function is preserved and above which it collapses; this is consistent with the clinically observed plateau-then-cliff pattern of dementia. The framework unifies cognitive reserve with channel coding theory, accounts for individual heterogeneity in disease onset, and generates falsifiable predictions that link information-theoretic redundancy measures to time-to-clinical-conversion.

## Introduction

Among individuals who accumulate comparable levels of amyloid-*β* plaques and hyperphosphorylated tau, clinical trajectories diverge sharply: some develop dementia within years of pathology onset, while others remain cognitively intact for decades (1, 2). The construct invoked to explain this divergence, *cognitive reserve*, is one of the most cited ideas in clinical neuroscience, with Stern’s foundational papers alone exceeding 10,000 citations in aggregate. Yet after roughly thirty years, cognitive reserve remains descriptive rather than mechanistic. It is operationalised through epidemiological proxies (education, occupational complexity, head circumference), it has no predictive theory that derives individual-level thresholds from information-theoretic quantities, and clinical onset is recognised only retrospectively (3).

In a seemingly unrelated development, Zheng and Meister recently quantified what they call “the largest unexplained number in brain science”: sensory neurons transmit information at a collective rate of order 10^9^ bits/s, while behavioral output proceeds at roughly 10 bits/s (4). The nine-order-of-magnitude gap has been framed as a puzzle about the serial nature of conscious processing (4), and subsequently debated: Sauerbrei and Pruszynski argued that unconscious sensorimotor processing substantially exceeds 10 bits/s and characterised the figure as a lower bound restricted to conscious cognition (5). Neither side, however, has considered what functional role this gap might serve in the context of disease.

We propose a synthesis: the 10^9^*/*10 gap is not merely a puzzle about consciousness; it can be understood, in substantial part, as *error-correcting redundancy* in the Shannon coding-theoretic sense. The vast number of neurons and circuits that mediate between sensory streams and the ∼ 10 bits/s behavioral bottleneck can be interpreted as redundant symbols in a biological error-correcting code. Under this identification, neurodegeneration-driven neuronal loss maps directly to symbol erasure, and the critical threshold for cognitive decline becomes derivable from coding theory rather than fitted post hoc.

The framework bridges three literatures that have not previously been connected in this way. The 10^9^*/*10 bottleneck (4) has attracted dozens of citations since 2024, but none route it through disease mechanisms. Biological error-correction codes (6, 7) have been applied to the reliability of spatial computation in grid cells, but not to progressive neuronal loss from neurodegeneration. Cognitive reserve and threshold models (1, 2, 8) have been discussed for three decades without an information-theoretic substrate. Recent graph-theoretic approaches to “brain functional redundancy” (17, 18) have measured observed correlates of reserve in neuroimaging data, but lack a theoretical framework that specifies what level of redundancy is sufficient and when it will be exhausted.

In what follows, we derive the critical damage fraction *d*_*c*_ = 1 − *k/n* from Shannon’s channel coding theorem, evaluate it across three complementary channel models, and show that the resulting framework: (i) produces a sharp phase transition matching the clinically observed “cognitive cliff,” (ii) accounts for heterogeneity in disease onset through individual variation in the redundancy ratio *ρ* = *n/k*, (iii) generates falsifiable predictions that distinguish it from existing threshold models, and (iv) predicts differential vulnerability of cognitive versus motor circuits from their respective bandwidth requirements.

## The framework

### The channel model

Consider a neural population of *n* functional units (neurons, cortical columns, or functional circuits) that collectively encode and transmit information from sensory input to behavioral output. The behavioral channel requires a throughput of *k* bits per processing window to sustain cognitive function, where *k* ≈ 10 bits/s corresponds to the conscious-cognitive bottleneck of Zheng and Meister (4). (Formally, *k* and *n* are both counts per processing window of duration *T* : the behavioral requirement is *k* = 10*T* bits per window, and *n* is the number of coding units available in that window. The code rate *R* = *k/n* and the threshold *d*_*c*_ = 1 − *R* are independent of *T* .)

In a healthy brain, the effective code rate is *R* = *k/n* ≪ 1: the system is massively over-provisioned relative to the minimum required throughput. We interpret this over-provisioning as the redundancy ratio *ρ* = *n/k* = 1*/R*.

Neurodegeneration removes functional units progressively. Let *d* ∈ [0, 1] denote the fraction of units that have been lost. The question is: how does residual cognitive capacity depend on *d*?

### The binary erasure channel

The simplest model treats each neural unit as a symbol in a block code transmitted over a binary erasure channel (BEC). Neuron loss at fraction *d* means each symbol is independently erased with probability *ε* = *d*. For a random linear [*n, k*] code, the block failure probability is

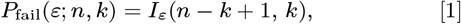

where *I*_*x*_(*a, b*) is the regularised incomplete beta function (22). By Shannon’s channel coding theorem (9), reliable decoding is possible when the code rate *R* is less than the channel capacity *C*_BEC_ = 1 − *ε*. Setting *R <* 1 − *ε* and solving for the critical damage fraction:

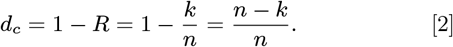

For biologically realistic parameters (*k* = 10, *n* = 1,000), *d*_*c*_ = 0.99: the code tolerates loss of 99% of its symbols before failure. The transition at *d*_*c*_ sharpens with increasing *n*, with width 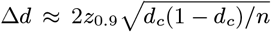, where *z*_0.9_ = 1.28 is the standard normal quantile.

### The Gaussian channel

A more biophysically grounded model treats each surviving neuron as contributing independent signal power *S* against fixed background noise *N*_0_. If the signals are pooled before noise is added (as in population vector decoding, where a downstream readout neuron sums over the population), total signal power scales linearly with surviving units: *S*_total_ = (1 − *d*) ·*n* ·*S*. (An alternative model treats each neuron as an independent parallel channel; this yields capacity linear in (1 − *d*) with no phase transition, and is discussed in the Limitations.) The Shannon–Hartley capacity is

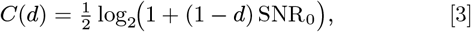

where SNR_0_ = *nS/N*_0_ is the baseline signal-to-noise ratio. Cognitive failure occurs when *C*(*d*) *< k*; this yields *d*_*c*_ = 1 − (2^2*k*^ − 1)*/*SNR_0_. For SNR_0_ = 10^8^ (consistent with 10^8^ neurons at unit SNR), *d*_*c*_ ≈ 0.99.

### Biological mapping

The quantity *n* is not the total number of neurons in the brain (∼ 86 billion (14)), but the number of functionally independent coding units in the circuit(s) subserving a given cognitive task. For a cortical column or functional circuit, *n* ∼ 10^3^–10^5^. The resulting *d*_*c*_ ∼ 0.99–0.9999 means local circuits tolerate 99–99.99% damage before failing, consistent with the clinical observation that substantial pathology precedes symptoms. For reference, normal aging produces only ∼ 10% neocortical neuron loss over the lifespan (19), well within the silent zone for any plausible *n*.

Critically, the redundancy interpretation differs from the naive duplication hypothesis that Zheng and Meister explicitly dismiss (4). They argue against “populations of essentially identical neurons” by citing the fruit fly (many cell types in only two copies) and the primate retina (one neuron per type per visual field point). Shannon-coded redundancy is categorically different: it distributes information across a population in structured, non-duplicative ways, as grid cells distribute spatial information via modular codes (6).

### Scope of the threshold

The critical fraction *d*_*c*_ = 1 − *k/n* is derived under independent symbol erasure, where each coding unit fails independently with probability *d*. This yields the sharpest possible threshold and is best understood as an *upper bound* on the damage a circuit can tolerate. In practice, neurodegeneration produces correlated damage: tau spreads along connected pathways, and vascular events affect contiguous tissue. Correlated erasure defeats coding redundancy more efficiently than independent erasure, because it can destroy an entire codeword’s worth of symbols in a single event. Under correlated damage, *d*_*c*_ shifts downward and the transition widens. The qualitative prediction (that a sharp threshold exists and depends on *ρ*) survives, but the specific numerical value *d*_*c*_ = 1 − *k/n* should be interpreted as a ceiling, not an exact prediction. Extending the framework to burst-erasure or spatially correlated failure models is a natural next step that would tighten the quantitative predictions.

### A worked example: entorhinal cortex

The entorhinal cortex layer II provides a concrete test of the framework. Gómez-Isla et al. (20) stereologically counted approximately 650,000 neurons in layer II of the entorhinal cortex in cognitively normal elderly controls. This region is critical for spatial and episodic memory and is among the earliest sites of tau pathology in Alzheimer’s disease.

The raw neuron count provides an upper bound on *n*, but individual neurons are not independent coding units: they share inputs, fire in correlated patterns, and serve multiple functions. The effective number of independent coding units *n*_eff_ is likely orders of magnitude smaller than the raw count. If we take *n*_eff_ = 1,000 (a conservative estimate corresponding to the number of functionally independent subcircuits or minicolumns), the framework gives *d*_*c*_ = 0.99 with transition width Δ*d* ≈ 0.008. If *n*_eff_ = 10,000, then *d*_*c*_ = 0.999 and Δ*d* ≈ 0.0008. The qualitative prediction (that *d*_*c*_ is close to 1, meaning substantial damage tolerance) holds across the plausible range of *n*_eff_.

Gómez-Isla et al. found that patients with very mild Alzheimer’s disease (CDR = 0.5) had already lost approximately 60% of layer II entorhinal neurons relative to controls. With *d*_*c*_ ≥ 0.99, a global damage fraction of *d* = 0.6 remains below the failure threshold, consistent with the observation that many individuals with substantial entorhinal neuron loss are still cognitively intact.

Importantly, 60% global loss does not imply uniform 60% loss in every local subcircuit. Tau pathology spreads along anatomically connected pathways and may devastate specific local circuits while sparing others. The theory predicts that spatial memory fails when *local* circuit damage in the critical subcircuit crosses *d*_*c*_, not when the population-level neuron count crosses an arbitrary threshold. This distinction between global and local damage fraction is testable with laminar-resolution post-mortem stereology and may explain why two patients with identical global entorhinal atrophy on MRI can have markedly different cognitive outcomes.

### Relationship to percolation theory

The coding-theoretic threshold *d*_*c*_ = 1 − *k/n* is information-theoretic: it depends on the ratio of required throughput to available coding units. A complementary threshold arises from percolation theory on random graphs. In an Erdős–Rényi network with mean degree *c*, random node removal destroys the giant connected component at a percolation threshold 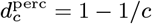 (16, 23). This is a graph-theoretic, not information-theoretic, quantity: it depends on network topology rather than information content.

The two frameworks make overlapping but distinguishable predictions. For brain networks where mean degree is high (*c* ∼ 10–100), both predict *d*_*c*_ near 1, and in the Erdős–Rényi simulation of Section 3, the percolation and coding thresholds are numerically similar. But they diverge for heterogeneous networks. A hub-and-spoke architecture has a lower percolation threshold than a homogeneous random network with the same mean degree, because targeted removal of hubs fragments connectivity disproportionately (16). The coding threshold, by contrast, depends only on the total number of functioning coding units regardless of their connectivity pattern.

This divergence has direct experimental consequences. In Alzheimer’s disease, tau spreads preferentially along highly connected hub regions (18). If damage is topology-dependent, percolation theory predicts faster network fragmentation than the coding-theoretic threshold would suggest, because hub removal is more destructive than random removal. Conversely, if the brain compensates by rerouting information through surviving pathways, the effective *n* is partially preserved and the coding threshold remains the binding constraint. Longitudinal connectomics data comparing hub-targeted versus diffuse atrophy patterns could distinguish these regimes and reveal whether cognitive decline in a given patient is connectivity-limited or capacity-limited.

## Analytical results

We evaluate the phase-transition prediction across five models of cognitive degradation under progressive damage.

### Binary erasure channel

Equation 1 produces sigmoid curves that sharpen with increasing *n* (Figure 2). For *n* = 5,000 and *k* = 10, the transition width is Δ*d* = 0.0016, effectively a step function. The linear degradation null (*P*_fail_ = *d*) dramatically overestimates failure at low damage and underestimates the cliff at *d*_*c*_. The critical threshold and transition width scale predictably with redundancy ratio *ρ* (Figure 3).

**Fig. 1.**
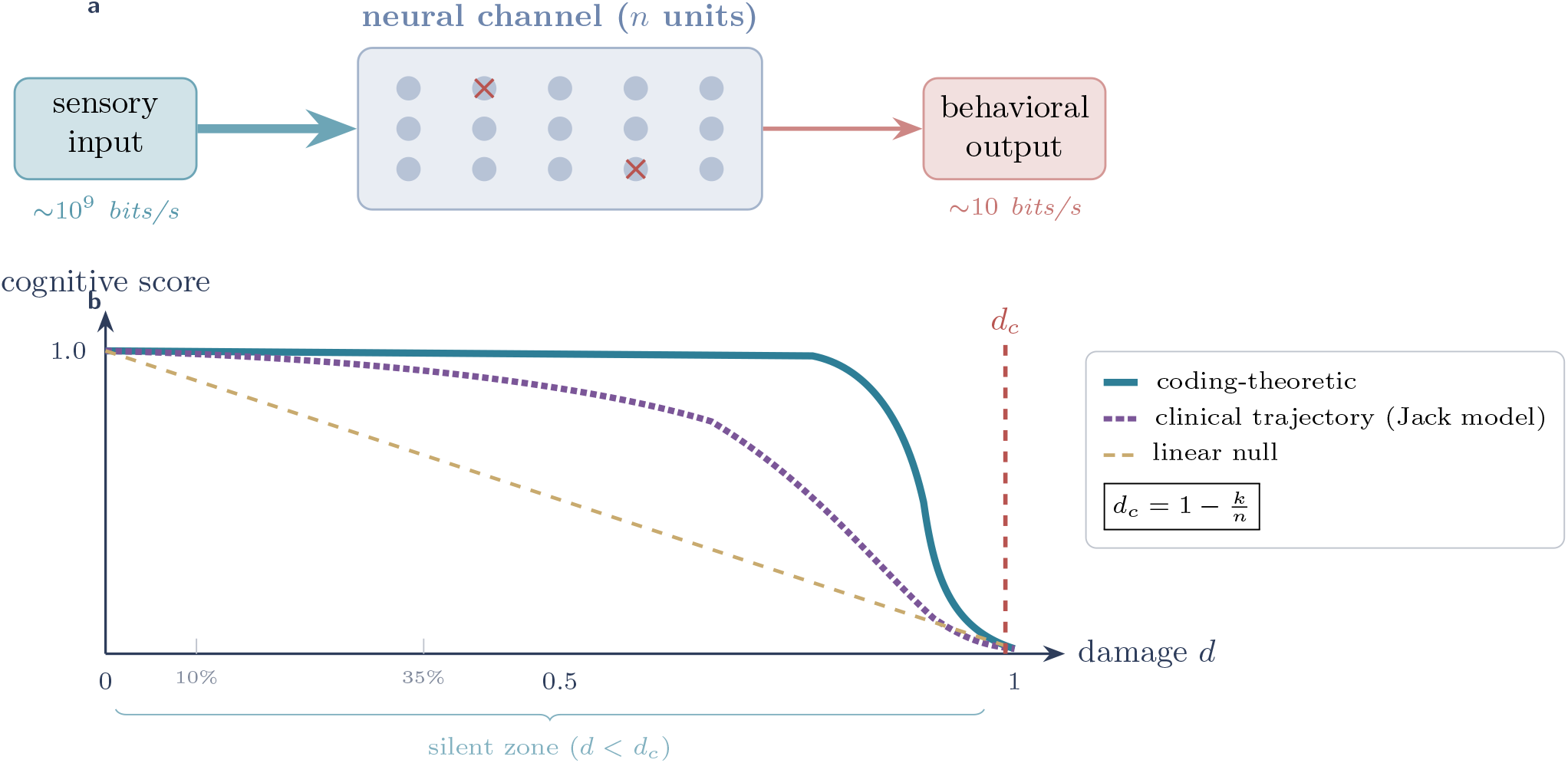
Conceptual framework. *Top:* sensory input (∼10^9^ bits/s) is encoded by *n* neural coding units and decoded to behavioral output (∼10 bits/s). Red crosses indicate units lost to neurodegeneration. *Bottom:* cognitive score as a function of cumulative damage fraction *d*. Three models are compared: the coding-theoretic threshold (teal solid), which predicts a long silent zone followed by a sharp cliff at *d*_*c*_ = 1 − *k/n*; the clinical trajectory as modeled by the Jack biomarker cascade (24) (purple dotted), a sigmoid with a subtle preclinical slope and a ∼3-year acceleration phase; and the linear degradation null (gold dashed). Vertical lines mark normal aging (∼10% loss) and early AD (∼35% loss), both within the silent zone. The equation is the coding-theoretic threshold, representing the upper bound under independent damage.

**Fig. 2.**
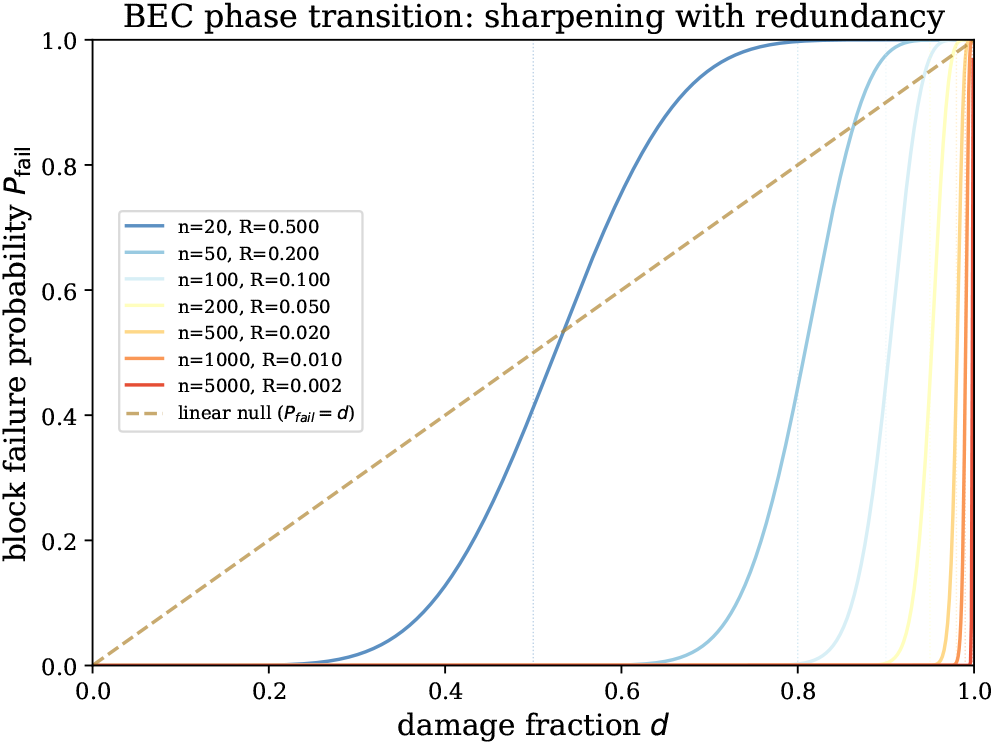
BEC phase transition. Block failure probability *P*_fail_ vs. damage fraction *d* for seven values of block length *n* (code rate *R* = *k/n*, with *k* = 10 fixed). Curves sharpen into a near-step-function as redundancy increases. Grey dashed line: linear degradation null. Vertical dotted lines mark *d*_*c*_ = 1 − *R* for each curve.

**Fig. 3.**
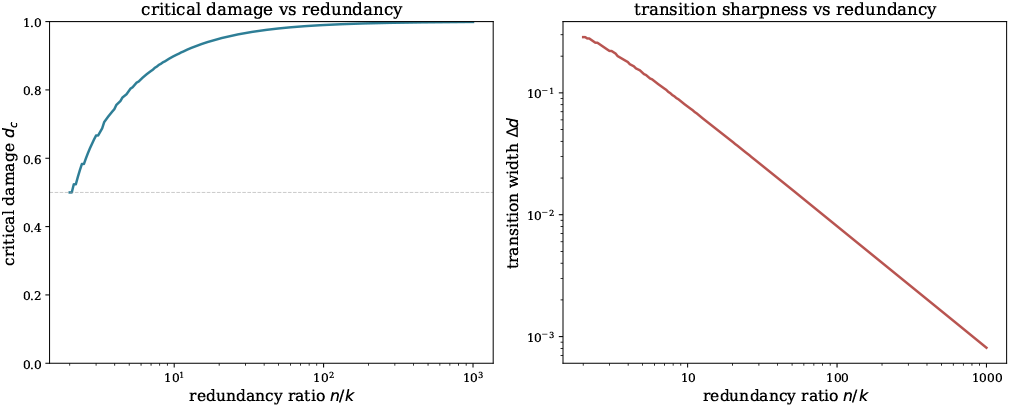
Scaling of *d*_*c*_ and transition width. *Left:* critical damage fraction *d*_*c*_ = 1 − 1*/ρ* as a function of redundancy ratio *ρ* = *n/k. Right:* transition width Δ*d* (the damage range over which *P*_fail_ goes from 0.1 to 0.9) decreases as 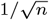, and produces an increasingly sharp cliff with larger circuits.

### Gaussian channel

The capacity waterfall (Equation 3) shows qualitatively identical behavior: a long plateau where *C*(*d*) ≫ *k*, followed by a steep drop when capacity approaches the task requirement (Figure 4).

**Fig. 4.**
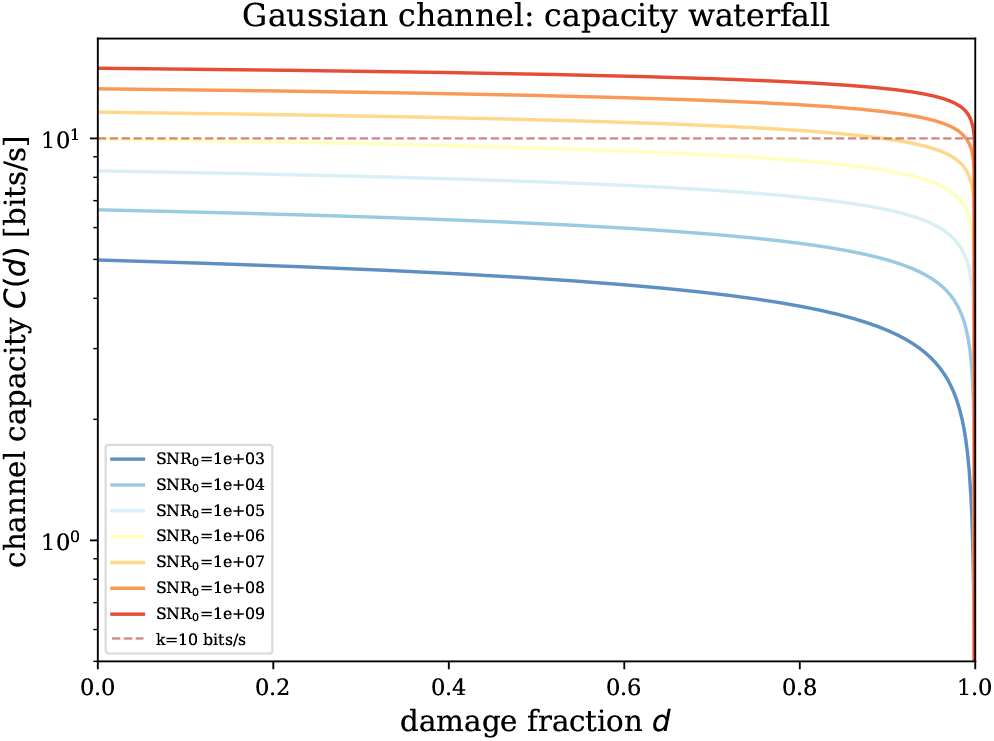
Gaussian channel capacity waterfall. Channel capacity *C*(*d*) vs. damage for seven baseline SNR values. Horizontal red line: task requirement *k* = 10 bits/s. Intersections mark *d*_*c*_ for each SNR. For SNR_0_ ≥ 10^7^, capacity remains far above the task requirement until damage approaches *d*_*c*_, then drops steeply.

### Erdős–Rényi percolation

As a bridge to network neuroscience, we model the brain as a random graph with mean degree *c*, where neuron loss removes nodes. The fraction of the giant component satisfies *g* = 1 − exp(− *c*(1 − *d*)*g*), with a percolation threshold at *d*_*c*_ = 1 − 1*/c*. This recovers a sharp transition driven by connectivity loss rather than information capacity, but with qualitatively similar cliff-like behavior.

### Linear null and Satz step

As comparators, we include the linear model (score = 1 − *d*, no threshold) and Satz’s (1993) step function (score = 1 for *d < d*_thresh_, 0 otherwise). The coding-theoretic models produce realistic intermediate sharpness between these extremes and, unlike the Satz step, *express* the threshold location in terms of physically interpretable quantities rather than fitting it as an unconstrained free parameter (Figure 6).

## Clinical predictions

### Individual heterogeneity from differential redundancy

The framework reduces the puzzle of cognitive reserve to a single parameter with physical interpretation: the redundancy ratio *ρ* = *n/k*. Here *k* ≈ 10 bits/s is constrained by the Zheng– Meister estimate, and *n* (the number of effective coding units in a given circuit) is a free parameter that must be estimated from neural data for each individual and each cognitive domain. Two individuals experiencing identical pathology rates (*α* damage/year) but differing in redundancy (*n*_*A*_ *> n*_*B*_, hence *d*_*c,A*_ *> d*_*c,B*_) will diverge in onset age by (*d*_*c,A*_ − *d*_*c,B*_)*/α* years (Figure 5). The population-level distribution of onset ages arises naturally from variance in *ρ*, complementing variance in pathology rate.

**Fig. 5.**
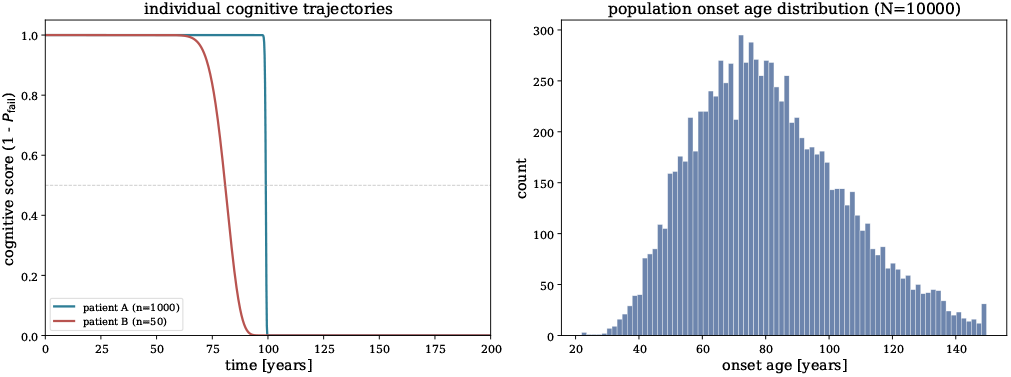
Clinical heterogeneity from differential reserve. *Left:* cognitive trajectories for two patients with identical damage rate (*α* = 0.01/year) but different redundancy (*n*_*A*_ = 1,000 vs. *n*_*B*_ = 50, *k* = 10). Both show a cliff, but shifted in time. *Right:* population-level onset age distribution (*N* = 10,000) arising from lognormal variation in *R* and *α*. Variance in onset reflects variance in reserve (*ρ*), not pathology rate.

**Fig. 6.**
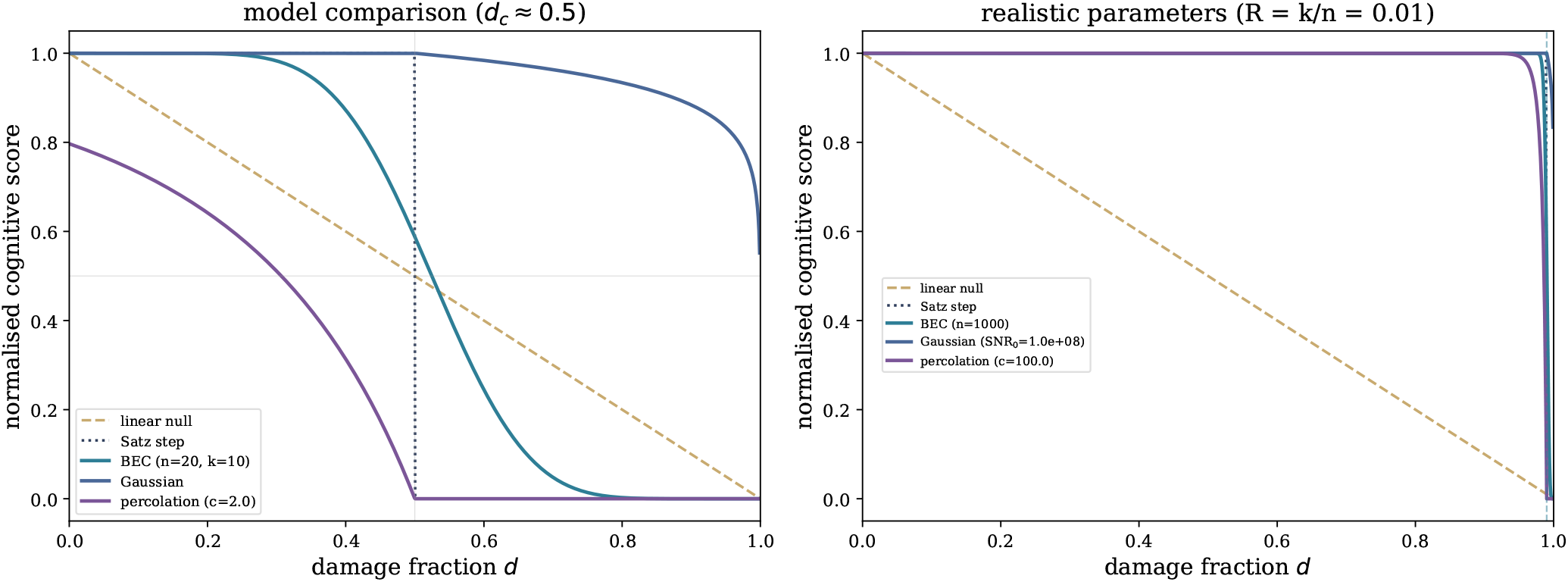
Model comparison. *Left:* five degradation models with matched *d*_*c*_ ≈ 0.5. The linear null (grey dashed) shows no threshold. The Satz step (black dotted) is discontinuous and requires *d*_*c*_ as a free parameter. The BEC, Gaussian, and percolation models produce smooth but sharp transitions whose threshold location is *derived* from the redundancy ratio. *Right:* realistic parameters (*R* = 0.01, which corresponds to a circuit of *n* = 1,000 coding units at *k* = 10 bits/s). All threshold models agree on *d*_*c*_ ≈ 0.99; only the coding-theoretic models express this value in terms of physically interpretable quantities.

This reframes Stern’s cognitive reserve as a quantitative, measurable object: *ρ* = *n/k*. Education, occupational complexity, and bilingualism (the standard epidemiological proxies) plausibly increase the effective *n* by expanding the repertoire of functionally independent neural coding units recruited for a given task. The framework also provides a natural account of why reserve is task-dependent: an individual may have high *ρ* for verbal tasks (many language-coding units) but low *ρ* for visuospatial tasks (fewer coding units), and this produces selective vulnerability that varies across cognitive domains within the same person.

### The cognitive cliff

Near *d*_*c*_, a small increment in damage produces a catastrophic drop in function. This is consistent with the “cognitive cliff” trajectory observed in a substantial subset of Alzheimer’s patients, where cognitive performance remains stable for years before accelerating sharply in the 2–3 years preceding diagnosis (12, 13). Not all patients show this trajectory (some decline gradually throughout), but the plateau-then-cliff pattern is well documented in longitudinal cohort studies and is the trajectory most difficult to explain under linear degradation models. The framework provides a quantitative account: the cliff can be understood as a coding-theoretic phase transition whose sharpness scales as 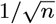.

The sharpness of the transition has a further clinical implication. In a circuit with *n* = 10,000 coding units, the transition width under independent erasure is Δ*d* ≈ 0.0008: at a damage rate of 1% per year, this corresponds to less than one month. In practice, correlated damage, compensatory plasticity, and measurement noise all broaden the transition, and the observed acceleration phase in longitudinal studies typically spans 1–3 years. This is consistent with the clinical observation that caregivers often describe cognitive decline as seemingly “sudden” even when biomarker evidence reveals pathology accumulating over decades. The framework predicts that the apparent speed of decline is not a property of the disease process itself but of the coding architecture: larger circuits (higher *n*) produce sharper cliffs. This also predicts that interventions arriving after *d*_*c*_ will have negligible effect (the code has already failed), while interventions arriving just before *d*_*c*_ could delay onset substantially by preserving even a small number of additional coding units.

### Testable predictions

Four predictions distinguish this framework from prior reserve models:

#### (i) Redundancy predicts onset beyond atrophy

Information-theoretic redundancy estimated from resting-state fMRI mutual information structure (15) should predict time-to-clinical-conversion beyond what structural atrophy alone predicts.

#### (ii) Task-specific vulnerability

Tasks with lower *k* (simpler output requirements) should tolerate more damage within a given circuit. The clinical ordering of cognitive decline in any disease depends on two interacting factors: which circuits are damaged first (a pathology question, governed by tau tropism, vascular territory, or genetic vulnerability) and which circuits have lower reserve (a bandwidth question, governed by *d*_*c*_ = 1 − *k/n*). In Alzheimer’s disease, Braak staging shows that pathology begins in entorhinal cortex and spreads through limbic to neocortical regions. Within each affected circuit, the framework predicts that higher-bandwidth functions (requiring more bits/s of behavioral output) will fail at lower damage fractions than simpler functions. This is testable by comparing the damage threshold for complex tasks (planning, multi-step reasoning) versus simple tasks (recognition, orientation) within the same brain region.

The framework also predicts that diseases targeting different circuits should produce different symptom orderings. In frontotemporal dementia, where neuronal loss is concentrated in prefrontal and anterior temporal circuits, behavioral symptoms (disinhibition, apathy, loss of social cognition) should appear before episodic memory deficits, which is precisely the clinical presentation that distinguishes behavioral-variant FTD from Alzheimer’s disease.

#### (iii) Complementarity with fluid biomarkers

Plasma p-tau217 indicates the presence of Alzheimer’s pathology; *ρ* governs *when* that pathology becomes clinically manifest. The combination of p-tau217 and *ρ* should outperform either alone for individualized prognosis.

#### (iv) Motor reserve is lower than cognitive reserve

The Sauerbrei–Pruszynski observation that motor control operates at higher bandwidth (*k*_motor_ ≫ 10 bits/s) directly implies *d*_*c*,motor_ *< d*_*c*,cognitive_ per Equation 2. Motor circuits, transmitting more information per unit time, have less redundancy per bit and thus lower damage tolerance. This predicts that motor symptoms should appear late in diseases that primarily damage cognitive circuits (Alzheimer’s) but early in diseases that primarily damage motor circuits (Parkinson’s, ALS). In Parkinson’s disease, the most affected subregion of the substantia nigra has lost ∼ 68% of its neurons by the time motor symptoms emerge (21), consistent with a motor channel operating at higher *k* with correspondingly lower but still substantial reserve. The asymmetry between cognitive and motor reserve, often noted clinically but rarely explained mechanistically, is consistent with the framework given the premise that motor bandwidth exceeds cognitive bandwidth for circuits of comparable size.

## Discussion

### Relationship to existing frameworks

Stern’s cognitive reserve (1, 2) is recovered as a special case: individuals with higher *ρ* have higher *d*_*c*_ and tolerate more pathology. The construct is no longer a descriptive label but a derived quantity with a physical interpretation.

Barrett, Deneve, and Machens (11) identified a “recovery boundary” for neuron loss using efficient coding theory and excitatory–inhibitory balance dynamics. Their framework describes the *circuit-level mechanism* that implements compensation; ours describes the *information-theoretic constraint* that determines when compensation must fail. The two are complementary layers of the same picture.

Cao (10) independently arrived at error-correcting code bounds for cognitive reserve, but for a different quantity: the combinatorial storage capacity of concept cells in the medial temporal lobe. Cao’s framework answers “how many concepts can be stored robustly?”; ours answers “how fast can information flow through the conscious bottleneck under progressive damage?” Together, these suggest the brain employs coding-theoretic redundancy at both the storage and transmission levels.

Simons, Levin, and Dichgans (12) proposed a “tipping points in neurodegeneration” framework using the dynamical systems metaphor of a ball rolling in a potential well. Our *d*_*c*_ provides a principled estimate of the well depth that their metaphor leaves unspecified. The dynamical systems and coding-theoretic pictures are not competing but describe different aspects of the same phenomenon: the dynamical framework captures the time-evolution of the system near the critical point, while the coding framework specifies where that critical point lies and how it depends on circuit architecture. Cotton et al. (13) extended the tipping point idea to cellular-resolution data and showed that individual neurons undergo abrupt transcriptomic state changes. Their cellular tipping points may correspond to the local coding-unit failures that accumulate toward the circuit-level *d*_*c*_ in our framework.

### The Sauerbrei–Pruszynski objection as a prediction

Sauerbrei and Pruszynski (5) argued that the 10 bits/s limit applies to conscious cognition but not to unconscious motor control, which employs substantially more bandwidth. Rather than undermining the framework, this objection generates a prediction. Motor control systems, operating at higher *k*, have lower *d*_*c*_ per Equation 2, meaning less reserve. This is consistent with clinical observations: in Alzheimer’s disease, motor function is typically preserved longer than higher cognition; in Parkinson’s disease, where damage targets the motor channel directly, motor symptoms appear first.

### Relationship to network redundancy measures

Recent experimental work has begun to quantify “brain functional redundancy” using graph-theoretic and information-theoretic measures applied to neuroimaging data. Schwarz et al. (17) measured dynamic brain functional redundancy (defined as the availability of alternative functional pathways that can sustain task performance when primary pathways are compromised) and proposed it as a neural substrate of cognitive reserve. Ghanbari et al. (18) found that network redundancy, measured via graph-theoretic disjoint paths (Menger’s theorem) among brain regions, increases during mild cognitive impairment (MCI) relative to healthy controls and then decreases in clinical Alzheimer’s disease.

This biphasic pattern (rise then fall) is predicted by the coding-theoretic framework. In the early stages of pathology (*d* ≪ *d*_*c*_), compensatory mechanisms upregulate alternative pathways; this effectively increases the number of functionally independent coding units *n* and thereby raises the measured redundancy. This compensation phase corresponds to the “silent zone” of Figure 1, where error correction masks accumulating damage. As pathology progresses and compensatory capacity is exhausted, effective *n* decreases, measured redundancy falls, and the system approaches *d*_*c*_.

The graph-theoretic “redundancy” of Schwarz et al. and the network redundancy of Ghanbari et al. can thus be understood as measured proxies for the information-theoretic redundancy ratio *ρ* = *n/k* that determines *d*_*c*_ in our framework. The coding-theoretic model provides the quantitative substrate that their experimental measures were missing: it specifies not only that more redundancy is protective, but provides a quantitative bound on how much damage a given level of redundancy can tolerate before the system fails. Connecting these observed graph-theoretic measures to the coding-theoretic *ρ* through longitudinal data would allow direct estimation of individual patients’ distance from their critical threshold.

### Limitations

The mapping from *n* to specific neural quantities is approximate. Real neural codes are not random linear codes over a binary erasure channel; they have structure, correlation, and heterogeneous connectivity. However, the phase-transition prediction is robust across channel models (BEC, Gaussian, percolation), which suggests the qualitative result (a sharp threshold at a redundancy-dependent *d*_*c*_) does not depend on the specific coding scheme. The Gaussian channel result (Equation 3) depends on coherent signal combining, which produces a logarithmic capacity curve and a sharp cliff. If neurons instead function as independent parallel channels, capacity degrades linearly with damage, reproducing the linear null. The BEC model does not depend on this assumption, and the phase transition in the BEC is the more fundamental result. The Gaussian model should be interpreted as illustrating how the cliff can arise in a continuous-valued setting when signal pooling occurs, not as a claim that all neural circuits pool signals coherently.

A further limitation is that we treat *n* as fixed for a given individual, whereas in reality the brain may dynamically recruit additional coding units through compensatory plasticity, to effectively increase *n* during early disease stages. This dynamic *n* would shift *d*_*c*_ upward in the short term while it potentially masks the approach to failure.

The framework is deliberately agnostic to the mechanism of damage (amyloid, tau, vascular, *α*-synuclein). This is both a strength (generality across neurodegenerative diseases) and a limitation (no specificity about which pathology drives *d* in a given patient). The implications of correlated damage for the threshold are discussed in the “Scope of the threshold” subsection above.

### Future directions

Four lines of work follow naturally. First, an experimental companion paper should estimate *ρ* from resting-state fMRI in a longitudinal cohort (e.g., ADNI) and test whether it predicts time-to-conversion beyond structural atrophy alone. The test is whether the information-theoretic redundancy measure adds predictive value over and above hippocampal volume and cortical thickness (the standard structural biomarkers). Second, a computational companion study will test the plateau-then-cliff prediction in a controlled setting by progressively pruning neural networks that have undergone grokking (a sudden generalisation phase transition) and measuring whether the generalising circuit collapses sharply at a critical damage threshold (Yin, in preparation). Third, the framework can be extended to specific neural architectures by replacing the idealized channel models with experimentally constrained network models that incorporate realistic connectivity patterns, cell-type heterogeneity, and spatially structured damage. Fourth, the distinction between coding-theoretic and percolation-theoretic thresholds developed here could be tested directly using longitudinal connectomics: patients whose decline is better predicted by hub connectivity loss than by total coding-unit loss are in the percolation-limited regime, while those whose decline tracks total unit loss are in the capacity-limited regime.

## Materials and Methods

All simulations were implemented in Python using NumPy and SciPy. The binary erasure channel failure probability (Eq. 1) was computed using `scipy.special.betainc.` The Gaussian channel capacity (Eq. 3) was evaluated analytically. The percolation giant component was found by solving *g* = 1 − exp(−*c*(1 − *d*)*g*) using `scipy.optimize.fsolve.` The population heterogeneity model sampled *N* = 10,000 individuals with lognormal-distributed code rate *R* (log-mean − 2.3, log-std 0.5, clipped to [0.001, 0.99]) and damage rate *α* (log-mean − 4.5, log-std 0.3); onset age was computed as (1 − *R*)*/α*. Total computation time: *<* 15 seconds on a consumer laptop.

## ACKNOWLEDGMENTS

D.Y. is supported by the Doctoral Training Programme in Medical Research (DTP-MR), University of Cambridge School of Clinical Medicine.

The author declares no competing interests.

All simulation code is available from the corresponding author upon request. No experimental datasets were generated or analysed in this study; all results are derived from analytical computations using publicly available software (Python, NumPy, SciPy). [4]

